# Predicting cell type-specific coverage profiles from DNA sequence

**DOI:** 10.1101/2025.06.10.658961

**Authors:** Johannes Linder, Han Yuan, David R. Kelley

## Abstract

Predicting expression profiles from RNA-seq experiments provides a powerful approach for universal sequence-based variant effect prediction, enabling researchers to score variants that affect total gene expression and relative isoform abundances. These models can be repurposed for new prediction tasks through transfer learning. However, current base models train primarily on bulk RNA-seq profiles derived from tissues and cell lines, overlooking the wealth of single-cell 3’-seq data that captures cell type-specific gene regulation. Here, we extend the capabilities of our recently developed *Borzoi* model by training on single-cell 3’-seq expression profiles from the Tabula Sapiens, Tabula Muris, and the Adult Brain Atlas, aggregated by cell type. This new model, *Borzoi Prime*, enables accurate variant interpretation across diverse cells, spanning erythrocytes to microglia. Training on 3’-seq profiles improves the model’s ability to predict cell- and tissue-specific alternative polyadenylation, even in the original bulk RNA-seq data. Through UTR-wide mutagenesis experiments of alternatively polyadenylated genes, we highlight determinants of cell type-specific 3’ UTR regulation learned by the model. This cell type-resolved approach opens new possibilities for understanding genetic variant effects via multiple layers of regulation in specific cellular contexts.

## Introduction

Understanding how genetic variants affect gene expression in specific cell types is crucial for interpreting disease-associated mutations and advancing precision medicine. Neural networks trained to predict regulatory function from DNA sequence have emerged as powerful tools for this purpose [1, 2]. By training on measurements from high-throughput sequencing assays of human cells and tissues, as well as those of closely related species, researchers have developed accurate models for diverse regulatory processes including enhancer activity, transcription initiation, splicing, mRNA degradation, and more [3, 4, 5, 6, 7, 8, 9, 10, 11, 12, 13, 14, 15, 16]. These models typically take genomic sequence as input and output either scalar or profile predictions representing specific regulatory functions.

We recently introduced *Borzoi*, a neural network that predicts RNA-seq expression profiles directly from DNA sequence, effectively unifying many of the aforementioned regulatory sequence models into a single network [17]. However, Borzoi was trained primarily on bulk sequencing experiments from tissues and immortalized cell lines [18, 19], preventing cell type-resolved predictions. This limitation is problematic because many trait-relevant variants exhibit cell type-specific effects.

Other research groups have addressed this gap through transfer learning. The Decima model predicts scalar gene expression values across diverse cell states [20], while the scooby model uses auxiliary cell embeddings to predict multiome profiles of continuous cell transitions [21]. However, no Borzoi-equivalent base model directly predicts expression profiles from diverse single-cell RNA-seq experiments at the cell type level.

Training sequence models on large collections of scRNA-seq profiles offers key advantages for downstream analyses. First, it enables direct cell type-specific variant scoring of total expression changes. Second, a base model trained on diverse cell types provides a stronger foundation for transfer learning applications (e.g. scooby). Third, most single-cell datasets use 3’-end sequencing protocols that capture alternative polyadenylation—a critical regulatory layer often disrupted in disease [22, 23]. Alternative polyadenylation varies across cell types, but standard RNA-seq models can only learn about it indirectly through subtle coverage pattern changes across terminal exons. In contrast, 3’-seq data directly captures cleavage site usage, with read coverage accumulating at each site proportional to its usage, enabling the model to learn sequence determinants of cell type-specific selection.

To address these limitations, we curated pseudo-bulk coverage profiles from diverse cell types across several large scRNA-seq atlases: Tabula Sapiens (human) [24], Tabula Muris (mouse) [25], Tabula Muris Senis (mouse) [26], and Adult Brain Atlas (human) [27]. We trained a new model, *Borzoi Prime*, on the combined single-cell data plus a subset of the original Borzoi training data. Borzoi Prime successfully learns to predict cell type-specific expression and effectively scores fine-mapped eQTLs in PBMCs and brain cell types. Training on 3’-seq profiles improves predictions of tissue- and cell type-specific alternative polyadenylation, even for bulk RNA-seq, and enables more accurate prioritization of polyadenylation QTLs. This work establishes a foundation for cell type-resolved variant interpretation and opens new avenues for understanding the cellular specificity of genetic effects in human traits.

## Results

### Predicting cell type-specific 3’-seq profiles from DNA sequence

Most scRNA-seq datasets are generated by 3’-end sequencing of polyadenylated mRNA transcripts [28, 29]. Consequently, sequencing reads accumulate at polyadenylation sites, resulting in 3’-biased expression profiles with distinct peaks upstream of 3’ cleavage sites within each gene. The relative peak magnitudes reflect site usage, while overall gene-level coverage reflects total expression [30, 31]. This signal complexity extends beyond simple polyadenylation patterns—intronic 3’ cleavage sites depend on upstream transcription start site (TSS) activity, creating interdependencies between transcription initiation and 3’-end processing. Additionally, cell- and tissue-specific splicing patterns influence compatible 3’ cleavage sites and may compete with the polyadenylation machinery for RNA processing. These interconnected regulatory layers make accurate scRNA-seq profile prediction challenging, requiring models that integrate information across entire transcript structures and their surrounding regulatory landscapes.

To develop a model capable of predicting diverse cell type-specific expression patterns, we curated scRNA-seq data from several large-scale atlases and aggregated alignments by cell type to generate pseudo-bulk profiles (Methods). These included the Adult Brain Atlas [27], Tabula Sapiens [24], and Tabula Muris (including Senis) [25, 26]. We combined these with auxiliary training data from the original Borzoi model, including CAGE, RNA-, ATAC-, and DNase-seq assays [18, 19, 32, 33], and optimized *Borzoi Prime* jointly on all datasets (Figure 1A). To further clarify transcript structure boundaries, we increased prediction resolution from 32 bp to 16 bp, yielding modest improvements on downstream evaluations. We trained an ensemble of four model replicates with identical train/test splits. The original Borzoi training set includes ATAC-seq profiles for many cell types also present in the newly added scRNA-seq data, likely improving model generalization.

**Figure 1:**
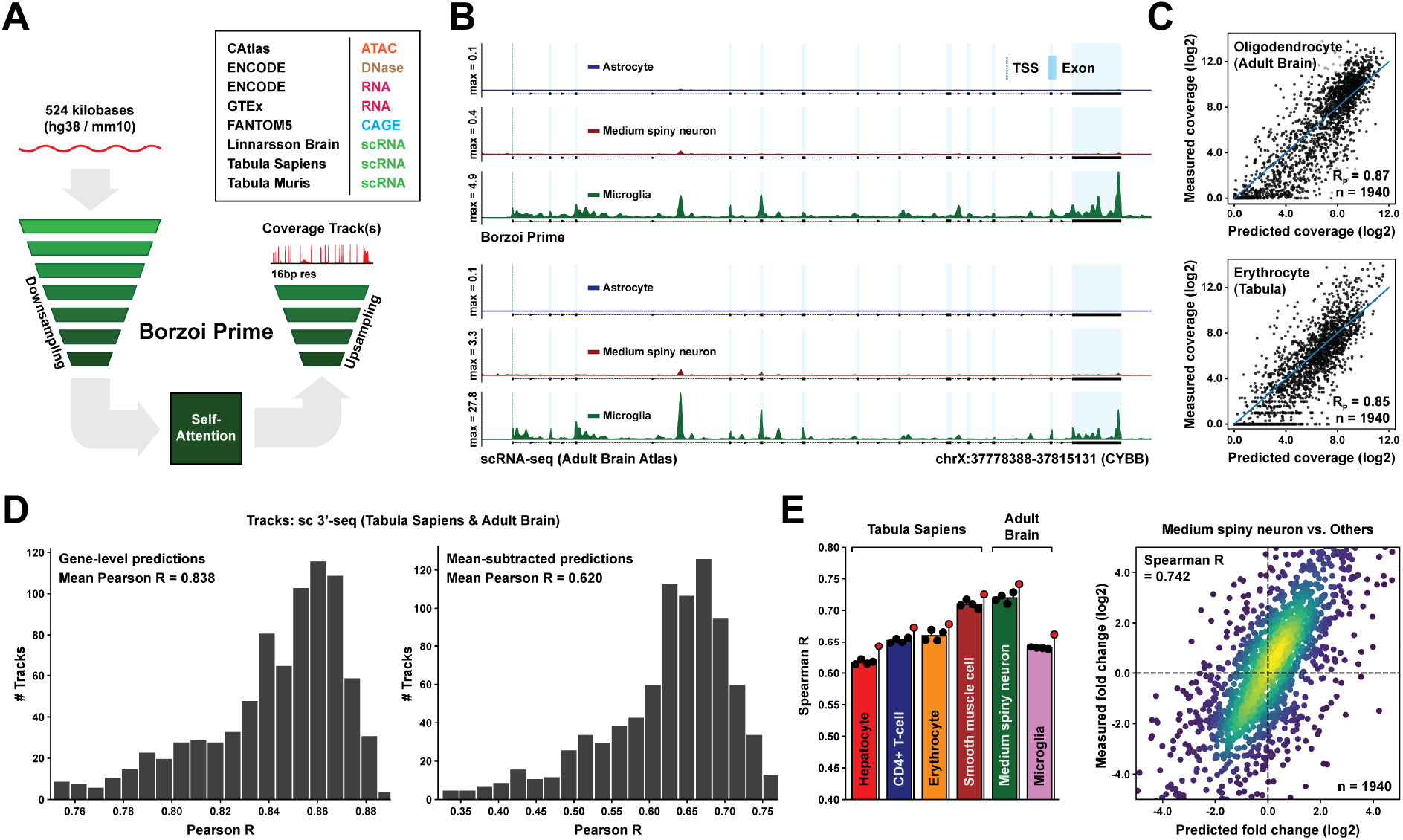
Borzoi Prime predicts cell type-specific 3’-seq coverage from DNA sequence. **(A)** Model architecture and training data overview. Borzoi Prime predicts coverage at 16 bp resolution for multiple epigenomic and transcriptomic assays, including 3’ scRNA-seq coverage tracks aggregated by cell type. **(B)** Example predictions for held-out gene *CYBB*, showing predicted (top) and measured (bottom) scRNA-seq coverage across brain cell types. **(C)** Scatter plots comparing predicted versus measured exon-aggregated scRNA-seq coverage for oligodendrocytes and erythrocytes on held-out genes. **(D)** Test set Pearson R distribution between predicted and measured exon-aggregated gene-level log-coverage (left) and quantile-normalized, mean-subtracted log-coverage (right) across scRNA-seq tracks. **(E)** Test set Spearman R predicting cell type-specific gene expression fold-changes relative to the average expression in five other cell types. Black dots represent replicate models, and red dots represent the model ensemble.

Despite the large distance between transcriptional regulation centered at promoters and transcript 3’ ends, Borzoi Prime achieved accurate predictions for 3’ RNA-seq profiles on held-out genes (Figure 1B-C). The model ensemble attained a mean Pearson R of 0.838 between predicted and measured gene-level log-normalized coverage values (Figure 1D, left) and accurately captured relative expression fold-changes between cell types (mean Pearson R = 0.620 across all tracks; Figure 1D, right). Across six well-characterized cell types, coverage fold-change Spearman R ranged between 0.65-0.74 (Figure 1E). Bulk RNA-seq performance was comparable to the original Borzoi model, with slight improvements in brain but reduced accuracy in blood samples (Supplementary Figure 1A-B). As expected, prediction accuracy for individual pseudo-bulk scRNA-seq tracks correlated strongly with sequencing depth (Supplementary Figure 1C).

### Sequence determinants of cell type-specific gene expression

To understand how Borzoi Prime identifies cell type-specific regulatory elements, we selected 1,000 genes with maximal differential expression across four Adult Brain Atlas cell types (Astrocyte, Medium spiny neuron, Microglia, and Oligodendrocyte) and another 1,000 differentially expressed genes across five Tabula Sapiens cell types (Erythrocyte, Hematopoietic stem cell, Hepatocyte, Fast muscle cell, and CD8+ T-cell). For each gene, we computed input-gated gradients of the exon-aggregated expression prediction [34], generating 1,000 saliency score vectors per cell type that quantify the contribution of all 524,288 input nucleotides to the model’s prediction. We observed that gradients from highly expressed genes sometimes exhibited saturation artifacts, prompting us to deploy an approximate integrated gradients approach for better calibration (Supplementary Figure 2A; Methods) [35]. However, for most genes, standard gradient saliencies were nearly identical to integrated gradients.

Figure 2A illustrates gradient interpretation for the *NGFR* gene, where integrated gradients highlight proximal and distal enhancers driving oligodendrocyte-specific expression (standard gradients shown in Supplementary Figure 2B). Zooming into these regulatory regions reveals a binding motif for SOX9/10, well-characterized oligodendrocyte transcription factors [36] (Figure 2B). Additional examples for other cell types and genes appear in Supplementary Figure 2C-D. To systematically identify cell type-specific regulatory motifs, we computed differential gradient saliencies by subtracting the average saliency across other cell types, then applied TF-MoDISco clustering [37] (Figure 2C; Supplementary Figure 3A). This analysis recovered known cell type-specific motifs, including RFX1/2/3 in neurons, IRF4/8 and CEBPA in microglia, and SOX9/10 in oligodendrocytes. Standard gradients yielded similar motif discoveries (Supplementary Figure 3B). To validate these computational predictions, we compared the aggregated cell type-specific saliency of each TF motif against measured expression fold-changes of the corresponding TF gene (averaging across TF subfamilies) between cell type pairs. This analysis revealed moderately high correlations for most cell type comparisons (Figure 2D, Supplementary Figure 3C; Spearman *R* = 0.25–0.91), supporting the biological relevance of the model’s learned regulatory logic.

**Figure 2:**
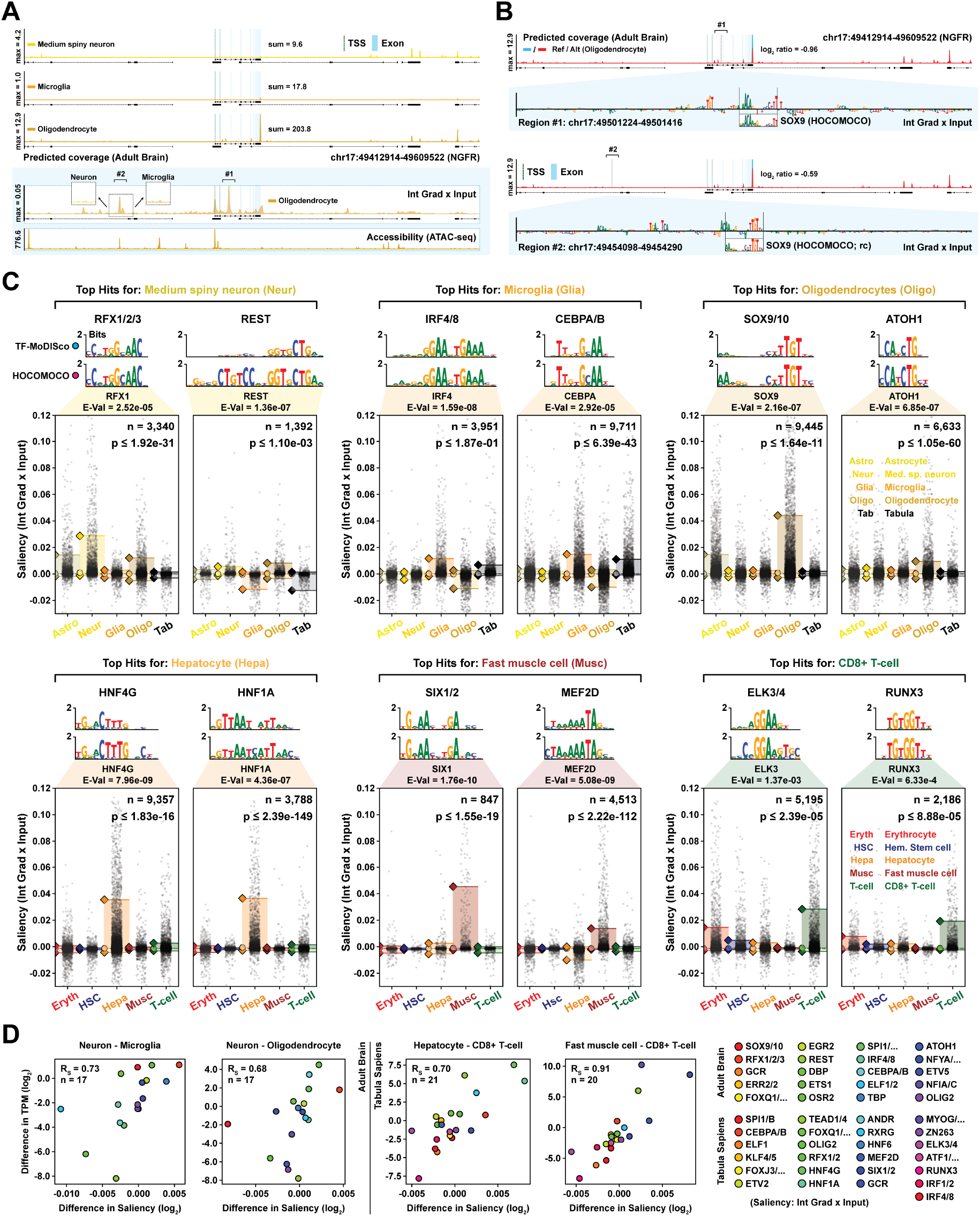
Model gradients reveal cell type-specific transcription factor motifs. **(A)** Predicted scRNA-seq coverage for gene *NGFR* aligned with integrated gradient attributions for exon-aggregated coverage and experimental ATAC-seq. **(B)** Predicted oligodendrocyte coverage when dinucleotide-shuffling proximal (top) or distal (bottom) putative enhancers, with gradient-based saliency scores displayed as sequence logos identifying SOX9/10 motifs. **(C)** Cell type-specific enrichment of selected TF motifs across six cell types. Swarm plots describe integrated gradient saliency distributions, with individual dots representing genomic motif instances. P-values from two-sided Wilcoxon tests compare the cell types with largest and second largest 95th percentile values. **(D)** The correlation between average motif saliency differences and corresponding TF gene expression differences (log-TPM, averaged across TF subfamily members) validates these motif inferences.

### Cell type-resolved prediction of expression-modifying genetic variants

Training on scRNA-seq profiles across diverse cell types enables Borzoi Prime to compute cell type-specific genetic variant effects on gene expression. Figure 3A demonstrates this capability using a fine-mapped causal eQTL SNP in microglia [38]. By comparing predicted expression profiles between reference and variant sequences and calculating the exon coverage fold change, the model correctly predicts that this variant increases *ADGRD1* expression. In-silico saturation mutagenesis (ISM) around the variant indicates gain of an ETS-like motif near other regulatory elements, including a putative MEF2D motif. The model predicts no effect in other brain cell types such as neurons or oligodendrocytes (Supplementary Figure 4A), demonstrating cell type-specific variant interpretation. Additional eQTL examples appear in Supplementary Figure 4B-C.

**Figure 3:**
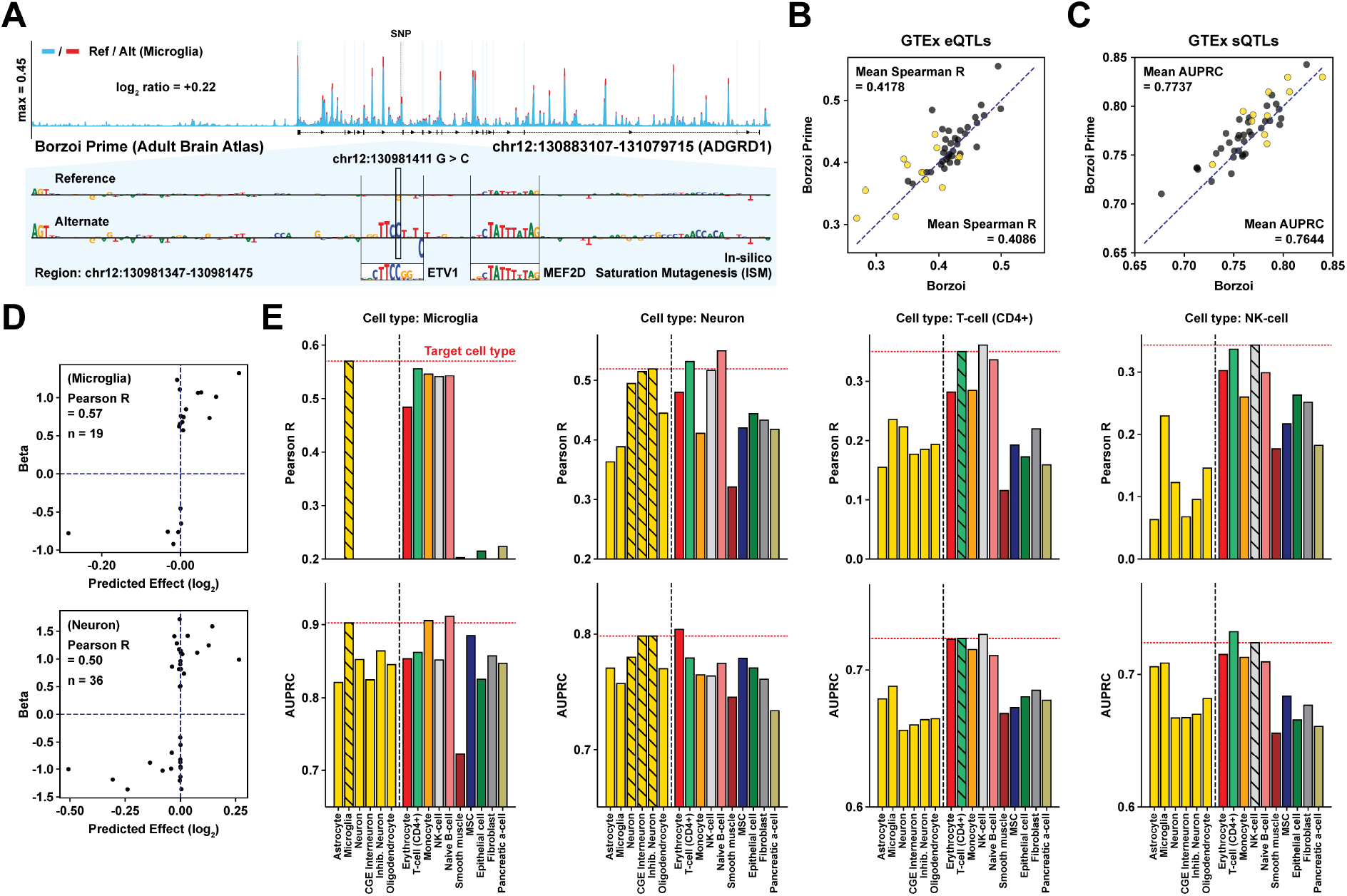
Cell type-specific prediction of genetic variant effects. **(A)** Example microglia eQTL prediction showing increased *ADGRD1* expression for the alternative (red) versus reference (blue) at SNP rs12821379, attributable to the creation of an ETS-like motif near a putative MEF2D motif matched to HOCOMOCO. **(B)** Spearman R between predicted and measured effect sizes for fine-mapped eQTLs comparing Borzoi Prime to the original Borzoi model across GTEx tissues. **(C)** Average precision for distinguishing fine-mapped sQTLs from matched negatives within 200 bp of splice junctions. **(D)** Predicted versus measured cell type-matched effect sizes for fine-mapped eQTLs in microglia and neurons. **(E)** Pearson R between predicted and measured scRNA-seq-derived effect sizes and average precision for eQTL classification across multiple cell types.

To assess whether training on single-cell 3’-seq data improved bulk tissue predictions, we evaluated Borzoi Prime on fine-mapped eQTL SNPs from GTEx [39] using scores derived from tissue-matched RNA-seq tracks. Performance marginally exceeded the original Borzoi model for gene-specific coefficient prediction (Figure 3B; mean Spearman *R* = 0.4178 versus 0.4086). We also calculated gene-agnostic variant effect scores based on all predicted tracks [17] and evaluated causal eQTL classification performance using random forest classifiers (8-fold cross-validation). Borzoi Prime achieved a mean AUROC of 0.7987 across tissues, an increase from Borzoi’s 0.7943 (Supplementary Figure 4D). Similar modest improvements appeared for bulk-tissue sQTL classification (Figure 3C; mean AUPRC = 0.7737 versus 0.7644 within 200 bp from an annotated splice junction). Supplementary Figure 4E illustrates improved prediction of splicing alterations caused by a well-characterized neurodegenerative disease risk variant (rs3115534 in *GBA1*) [40].

We next evaluated cell type-specific variant prediction by curating eQTL Catalog datasets [41, 42] for microglia [38], neurons [43], and scRNA PBMCs [44], constructing matched negative SNP sets for each cell type (Methods). We derived variant effect scores from predicted expression profiles using Adult Brain Atlas or Tabula Sapiens tracks and evaluated both classification performance and correlation with experimental effect sizes. To improve comparison across cell types, we incorporated track-specific pseudocounts to downweight fold changes in lowly expressed genes (Methods). Correlations between predicted and measured effect sizes ranged from 0.35 to 0.57 (Figure 3E). Variant scores derived from cell type-matched predictions consistently outperformed off-target cell types—e.g. microglia-derived scores showed highest correlation with microglia eQTL effects. This cell type specificity was also evident in eQTL classification tasks. Additional evaluations for other cell types appear in Supplementary Figure 4F.

In the supplement, we compared Borzoi Prime and scooby [21] for eQTL prediction across a subset of PBMC cell types (Supplementary Figure 4G). scooby achieved higher Spearman correlation with the measured effects, while Borzoi Prime achieved higher Pearson correlation. This indicates that the models have complementary strengths: scooby better captures rank-order relationships across the full range of effect sizes, while Borzoi Prime better captures linear relationships for variants with larger effects. Consistent with this, scooby showed fewer sign discordances for variants with small effect sizes.

### Predicting alternative polyadenylation across cell types

Finally, we evaluated whether Borzoi Prime learned to predict cell type-specific alternative polyadenylation from the scRNA-seq training data. By examining peaks near annotated polyadenylation sites in predicted 3’-seq profiles of held-out genes, we observed striking differences in site usage between cell types that matched measured profiles (Figure 4A, left). The model also gave consistent predictions for bulk RNA-seq profiles of matched tissues (Figure 4A, right). We observed a general trend of 3’ UTR lengthening in neurons and shortening in blood cell types such as erythrocytes, consistent with prior work [45, 46, 47].

**Figure 4:**
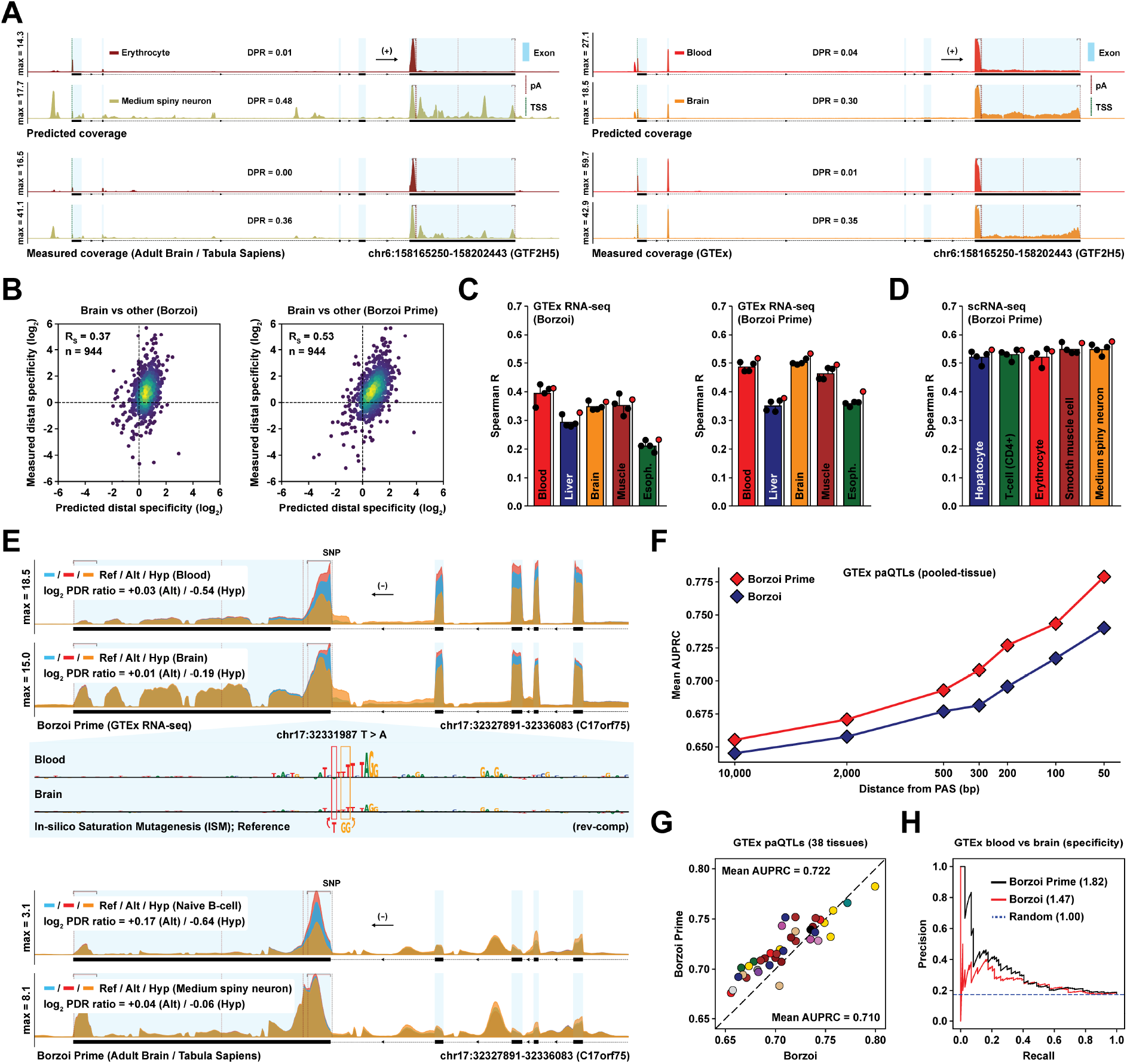
Predicting tissue- and cell type-specific alternative polyadenylation. **(A)** Predicted and measured coverage for held-out gene *GTF2H5* in (left) scRNA-seq blood and brain cell types and (right) bulk RNA-seq blood and brain tissues. DPR = distal-to-proximal polyadenylation site coverage ratio. **(B)** Predicted versus measured distal polyadenylation usage in brain tissue relative to the average of four other tissues for held-out test genes. **(C)** Spearman R for predicted versus measured distal polyadenylation usage ratios for five tissues relative to the other four. Black dots represent replicate models, and red dots represent the model ensemble. **(D)** Spearman R for predicting distal polyadenylation usage ratios across cell types using scRNA-seq tracks. **(E)** Predicted RNA-seq coverage in blood and brain for the reference (blue), paQTL variant rs1019152 (red), and hypothetical double-mutation AA*>*CC that weakens the polypyrimidine tract of gene *C17orf75* (orange). PDR = proximal-to-distal coverage ratio. **(F)** Average precision for Borzoi and Borzoi Prime when distinguishing fine-mapped paQTLs from matched negatives using variant scores derived from GTEx RNA-seq tracks in tissue-pooled analysis. **(G)** Average precision per GTEx tissue for distinguishing fine-mapped paQTLs from matched negatives within 10000 bp of a PAS. **(H)** Precision-recall curves for classifying tissue-specific paQTL effects in brain versus blood-specific or shared effects.

To understand sequence determinants driving differential polyadenylation, we performed in-silico saturation mutagenesis across 3’ UTRs of genes showing neuron-erythrocyte differences in isoform usage. Surprisingly, the 3’-most splice acceptor and neighboring regulatory elements were often the most salient features, alongside less prominent U-rich motifs near distal polyadenylation signals (Supplementary Figure 5A). Proximity between the 3’-most splice acceptor and proximal polyadenylation signal associated with predicted non-neuronal shortening, consistent with known interactions between splice acceptor efficiency and proximal cleavage [48, 49, 50, 51, 52]. Supporting this prediction, we found significant correlations between proximal PAS distance relative to the 3’-most splice acceptor and non-neuronal proximal PAS preferential usage in both GTEx and PolyASite 2.0 data [53] (Supplementary Figure 5B). Additional examples of brain-biased distal polyadenylation appear in Supplementary Figure 5C-E.

We next compared Borzoi Prime to the original Borzoi model for predicting distal-to-proximal coverage ratios across bulk RNA-seq tissues for held-out genes. Borzoi Prime was substantially more accurate (Figure 4B-C), suggesting successful transfer of 3’-seq regulatory features to bulk RNA prediction. The model also accurately predicted cell type-specific 3’ UTR lengthening from scRNA-seq tracks (Spearman *R* = 0.53–0.55 across five cell types; Figure 4D).

Given these improvements, we hypothesized that Borzoi Prime would better score polyadenylation QTLs (paQTLs). Figure 4E highlights a fine-mapped paQTL that strengthens the 3’-most splice acceptor polypyrimidine tract, increasing proximal 3’ cleavage (rs1019152), alongside a hypothetical variant predicted to decrease proximal cleavage. Consistent with our findings, the model predicted larger effects in blood and immune cells than brain, matching experimental observations that terminal exon effects are ∼2× larger in lymphoblastoid cell lines than cerebellum (as visualized in Elixir [54]). See Supplementary Figure 6A-D for additional examples. When classifying fine-mapped causal paQTLs from matched negatives, Borzoi Prime outperformed Borzoi (Figure 4F-G, Supplementary Figure 6E). The model also excelled at the challenging task of predicting tissue-specific paQTL effects, successfully distinguishing blood-specific from brain-specific variants based on predicted effect differences (Figure 4H, Supplementary Figure 6F; Methods). Supplementary Figure 6G provides additional examples of known pathogenic polyadenylation-altering variants.

## Discussion

We developed Borzoi Prime, a neural network that extends sequence-based gene expression prediction to cell type-specific contexts. By training on 3’ single-cell RNA-seq data from three large atlases—Tabula Sapiens [24], Tabula Muris [25], and the Adult Brain Atlas [27]—alongside auxiliary epigenetic and transcriptional datasets, Borzoi Prime predicts cell type-specific 3’-seq expression profiles with high accuracy on held-out genes. The model accurately predicts both cell type-specific expression and 3’-processing patterns, enabling robust cell type-resolved variant interpretation, as validated through fine-mapped eQTL studies in neurons, microglia, and blood cells. Training on 3’-seq data improved predictions of tissue-specific polyadenylation QTLs compared to the original Borzoi [41].

Beyond variant prediction, Borzoi Prime provides insights into cell type-specific regulatory mechanisms. Through gradient-based interpretation, we recovered known TF motifs for each cell type. Our analysis of alternatively polyadenylated genes in neurons revealed an interesting mechanistic insight: the model learned that proximity between the 3’-most splice acceptor and proximal polyadenylation sites drives tissue-specific UTR shortening, consistent with known competition between splicing and polyadenylation machinery [48].

Borzoi Prime addresses a critical gap in regulatory sequence models by providing cell type-resolved expression profile predictions. The benefits complement other recent cell tools: scooby predicts expression across continuous cell states using multiome data [21], and Decima focuses on scalar gene expression values across harmonized studies [20]. In contrast, Borzoi Prime uniquely combines high-resolution profile-level predictions with diverse cell type coverage, enabling both isoform-level analysis and seamless integration with epigenomic datasets. This combination of profile-level detail and cell type diversity positions the model as a powerful foundation for transfer learning applications requiring cell type-specific regulatory understanding.

## Methods

### Training data

We curated 3’ scRNA-seq data from: Tabula Sapiens [24], Tabula Muris [25], Tabula Muris Senis [26], and the Adult Brain Atlas [27]. For each dataset, we downloaded alignments for hg38 or mm10, split aligned reads based on cell type cluster assignments (using cell barcodes), and aggregated pseudo-bulk coverage tracks for each annotated cluster. In total, we obtained 1,702 stranded scRNA-seq coverage tracks for 851 distinct clusters.

We included these data in Borzoi Prime’s training set, along with most original Borzoi [17] training data: human and mouse CAGE data from FANTOM5 [32, 55], RNA-seq and DNase-seq data from GTEx and ENCODE [18, 56, 19] and ATAC-seq data from CATlas [57, 33]. Finally, we included tissue-pooled polyadenylation peak tracks from PolyADB V3 [58] and PolyASite 2.0 [53]. We excluded ENCODE ChIP-seq data to limit the growing number of output tracks, hypothesizing that DNase/ATAC assays sufficiently captured epigenomic signals. The collection includes cell type-specific coverage tracks present in the ATAC-seq data, including for microglia, oligodendrocyte, and more.

We applied similar transformations to the binned and aggregated coverage values as in the original Borzoi paper, with minor improvements. Unlike Borzoi, we add +1 to coverage values before the exponential squash transform to mitigate inflating values less than 1. The squash formula is given below:

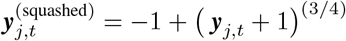

We similarly add +1 before soft-clipping values above a certain threshold, to refrain from inflating residuals less than 1. The updated soft-clip formula is given below, where *c* denotes the threshold (*c* = 64 for RNA-seq and scRNA-seq):

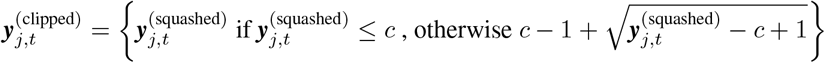

As in the original Borzoi dataset construction, we fragmented the human and mouse genome into 8 data folds (keeping orthologous sequences in the same fold) and trained four model replicates with identical train/test splits to the original Borzoi. We used a modified human reference genome for input sequences, where the allele with maximum frequency in gnomAD v3.1 [59] was substituted at each position.

### Model architecture and training

The model follows the same architecture as Borzoi, consisting of an initial convolutional tower with max-pooling layers, a stack of self-attention layers, and a stack of convolutional and upsampling layers with U-net connections. We added one upsampling layer to model coverage at 16 bp resolution given a 524 kb input sequence, fitting with batch size 2× across 2 NVIDIA A100 GPUs. Each model replicate was trained using the Adam optimizer until bin-level validation Pearson R saturated.

### Input sequence attribution

Given a summary statistic *u*(***x***) ∈ ℝ calculated from one of Borzoi Prime’s predicted coverage tracks ***y*** = ***f*** (***x***) ∈ ℝ^*L*×*T*^ (where *L* is the output length and *T* is the number of tracks), we computed *contribution scores* of *u*(***x***) using four techniques. The first three—input-gated gradient saliency (‘Grad x Input’), in-silico saturation mutagenesis (‘ISM’), and window-shuffled ISM (‘ISM Shuffle’)—were used exactly as described in the Borzoi publication. We also re-used previously defined summary statistics, including log-sum of exon coverage and log-coverage ratios. However, as described in the results section, we observed that some highly expressed genes exhibited saturated gradients, resulting in contribution scores that were not proportional to the magnitude of the prediction. This phenomenon was prevalent for some, but far from all, highly expressed genes. To derive better calibrated contribution scores for this small set of genes, we employed an approximate ‘Integrated Gradients’ method [35] that stochastically breaks saturation by introducing varying amounts of random noncoding mutations and averaging gradients across mutants. The approach is similar to Expected Gradients and related methods [60, 61, 62]. The details are given below:

We first define a sequence of noise levels *σ*_*t*_ ∈ [0, 1], corresponding to the fraction of noncoding positions to randomly mutate for each gradient sample. In Supplementary Figure 2A, we use noise levels *σ*_1_, *σ*_2_, … = 0.00, 0.02, 0.05, 0.10, 0.15, 0.20, 0.25, 0.30, 0.35. However, for the large-scale gradient analysis presented in Figure 2, we limited the noise levels to *σ*_1_, *σ*_2_, … = 0.00, 0.02, 0.05, which often sufficed to break saturation effects. Next, for a given noise level *σ*_*t*_, we create *K* sequence mutants 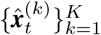, where each mutant 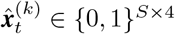 has a fraction *σ*_*t*_ of its non-exon-overlapping regions randomly mutated relative to wild type sequence ***x*** ∈ {0, 1}^*S*×4^.

For each sequence mutant, compute the gradient saliency as follows: 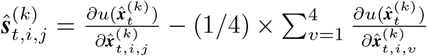

Finally, calculate the total saliency as the unweighted mean across samples: 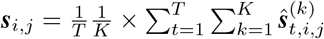

Note 1: For expression gradients, we use *u*(***x***) = log (*C* + (1*/*|ℋ |) × ∑_*h* ∈ ℋ_ ∑_*b* ∈ ℬ_ ***y***_*b,h*_), where ℋ is the set of target indices, ℬ is the set of exon-overlapping bin positions, and *C* ∈ ℝ is an optional pseudo count.

Note 2: The analysis presented in Figure 2 was based on the approximate integrated gradients method described above. However, vanilla input-gated gradients produced well-calibrated scores for most genes, and produced near-identical TF-MoDISco results as the version with integrated gradients (Supplementary Figure 3B).

### Cell type-specific motif clustering

We selected 800 genes with highest variance-to-mean ratio (VMR) across four top-level cell clusters from the Adult Brain Atlas (Astrocyte, Medium spiny neuron, Microglia, and Oligodendrocyte) plus 200 genes with highest brain/non-brain fold change in GTEx. We also selected 1000 genes with highest VMR across five Tabula Sapiens cell types (Erythrocyte, Hematopoietic stem cell, Hepatocyte, Fast muscle cell, and CD8+ T-cell). We computed input gradients of exon-aggregated log-transformed gene expression predictions (average across sub-cluster coverage tracks belonging to the same major cell type), subtracted the average gradient of other cell types, and ran TF-MoDISco on the target cell type saliencies [37]. We pooled MoDISco results across cell types by matching the highest-ranking TF of each motif cluster. We annotated TFs using the Tomtom MEME suite [63] to query the HOCOMOCO v11 database [64]. When comparing TF motif saliencies against corresponding TF gene TPM values (Figure 2D, Supplementary Figure 3C), we manually inverted the saliency sign of a small number of TFs with known repressive function—REST, OLIG2, MyoD.

### Expression, splicing, and polyadenylation QTL benchmarks

We used the expression, splicing, and polyadenylation QTL datasets from GTEx v8 curated in the original Borzoi publication, including the same set of distance-matched negative (non-causal) SNPs. For expression QTLs (eQTLs), we compared Borzoi Prime to Borzoi at both gene-agnostic eQTL classification (based on L2 scores) and gene-specific eQTL coefficient prediction (based on logSED scores). For gene-agnostic eQTL classification, we derived L2 variant scores for all predicted tracks and fit a random forest classifier to distinguish between fine-mapped eQTLs and matched negatives based on all scores (8-fold cross-validation). For gene-specific coefficient prediction, logSED scores were derived from tissue-matched RNA-seq tracks. For splicing QTLs (sQTLs), we derived variant scores from the maximum normalized predicted difference in coverage. For polyadenylation QTLs (paQTLs), we derived scores from the maximum predicted 3’ site coverage ratio fold change. We refer to the original Borzoi publication for definitions of these variant scores [17]. For both splicing and polyadenylation variant effect prediction, we derived classifier scores from tissue-matched RNA-seq tracks only.

In addition to bulk-tissue QTL benchmarks, we curated cell type-resolved fine-mapped eQTL sets from the eQTL Catalogue [41, 42]. These included three studies: (1) bulk- and scRNA-seq of primary human microglial cells [38], (2) bulk RNA-seq of cultured neuronal cells [43], and (3) scRNA-seq of PBMCs from 982 donors [44]. We constructed negative SNP sets for each study, matching for TSS distance, cell type-specific expression, and allele nucleotides. For the neuronal eQTL set, we filtered fine-mapped SNPs for posterior inclusion probability (PIP) ≥0.9, resulting in 36 positives. For the microglia eQTL set, due to the relatively small sample size, we lowered the PIP threshold to 0.7, resulting in 19 positives. For the sc-eQTL study in PBMCs, we retained only cell types with sufficiently large numbers of fine-mapped SNPs with PIP ≥0.9 that we also had cell type-matched scRNA-seq tracks for in the training data. Specifically, we evaluated predictions for CD4+ T-cells (233 positives), NK cells (201 positives), CD4+ TCM T-cells (246 positives), CD8+ TEM T-cells (104 positives), Naive B-cells (50 positives), and Monocytes (25 positives). To compare against scooby [21], we downloaded pre-computed variant effect scores made by their fine-tuned OneK1K model and intersected them with Borzoi Prime predictions.

We evaluated Borzoi Prime cell type-resolved, gene-specific eQTL classification and coefficient prediction based on the logSED statistic, derived from coverage tracks corresponding to a single cell type. Since tracks for off-target cell types often predict low baseline exon coverage, we added a track-specific pseudocount to the predicted reference and alternate coverage sums before computing their log fold change. The pseudocount was defined as the 20th percentile of predicted expression across all genes. Due to small sample sizes of some eQTL sets, we used Pearson R rather than Spearman R to evaluate coefficient predictions.

Finally, to evaluate the model’s ability to identify tissue-specific paQTLs, we constructed a classification task from fine-mapped GTEx paQTLs. We defined tissue-specific positive paQTLs as variants with PIP ≥0.9 in the target tissue and PIP *<* 0.2 in non-target tissues, and used as negatives paQTLs with PIP ≥0.2 in non-target tissues. We classified tissue-specific paQTLs from these negatives based on the difference between predicted variant effects derived from the maximum 3’ log coverage in the target tissue versus the average effect in non-target tissues.

## Availability of data and software

The code repository for the original Borzoi model was used to train and evaluate Borzoi Prime (https://github.com/calico/borzoi). Pre-trained model weights and evaluation scripts are available at: https://github.com/calico/borzoi-paper/tree/main/extensions/prime. The processed Borzoi Prime training data are available for download at: gs://prime-paper/data/ (Google Cloud Storage). Fine-mapped GTEx QTL sets and matched negatives are available at gs://borzoi-paper/qtl/. Cell type-resolved eQTL sets were curated from the eQTL Catalogue (https://www.ebi.ac.uk/eqtl/), and are available in a processed format at gs://prime-paper/qtl/. Gene annotations were obtained from: https://www.gencodegenes.org/ (v41). Predictions for the scooby model were obtained from: https://github.com/gagneurlab/scooby_reproducibility.

The Genotype-Tissue Expression (GTEx) Project was supported by the Common Fund of the Office of the Director of the National Institutes of Health. Additional funds were provided by the NCI, NHGRI, NHLBI, NIDA, NIMH, and NINDS. The datasets used for the analyses described in this manuscript were obtained from dbGaP at http://www.ncbi.nlm.nih.gov/gap through dbGaP accession number phs000424.v9.p2.

## Author’s Contributions

Conceptualization: D.R.K.; Analysis: J.L., H.Y., D.R.K.; Writing: J.L., H.Y., D.R.K.

## Acknowledgements

We thank Michael Closser for helpful discussions and advice on enhancer-mediated gene regulation.

