## Supplementary Figure 1-6 for "Predicting cell type-specific coverage profiles from DNA sequence"

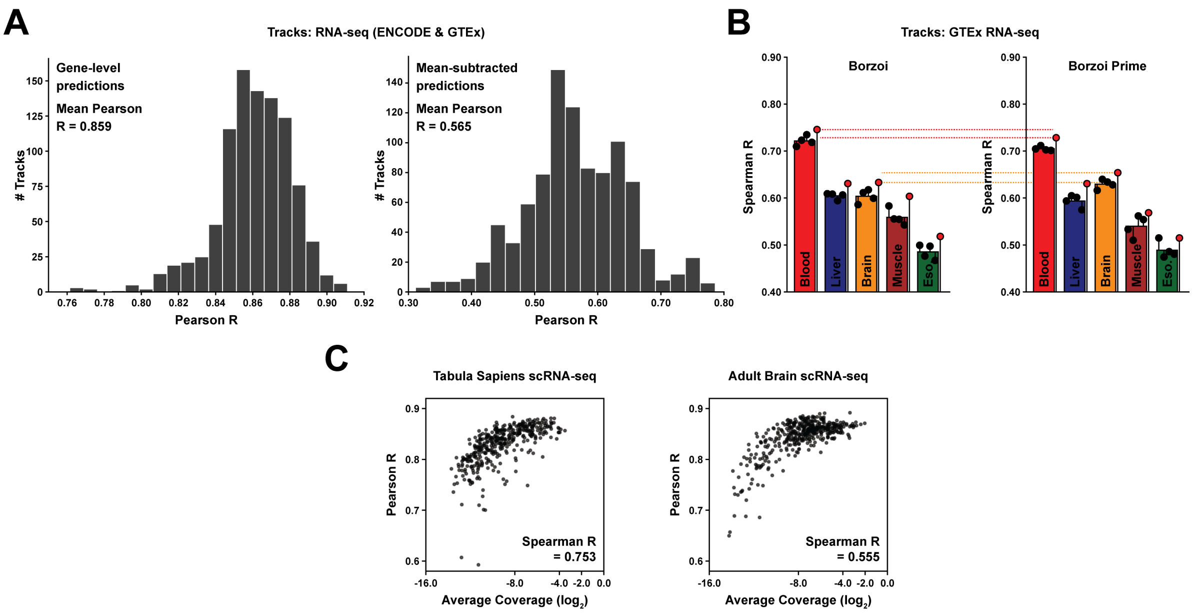


Supplementary Figure 1: **Related to Figure 1.** **(A)** Test set Pearson R distribution for bulk RNA-seq tracks when predicting gene-level, exon-aggregated log-coverage values (left) or quantile-normalized, mean-subtracted log-coverage values (right). **(b)** Test set Spearman R comparing predicted to measured gene-level fold changes across GTEx bulk RNA-seq tissues. The fold changes are calculated as the ratio of exon-aggregated coverage in one tissue relative to the average coverage in 4 other tissues. Shown are the results for Borzoi (left) and Borzoi Prime (right). **(C)** Comparison between average genome-wide sequencing coverage and held-out gene-level test performance for scRNA-seq tracks.

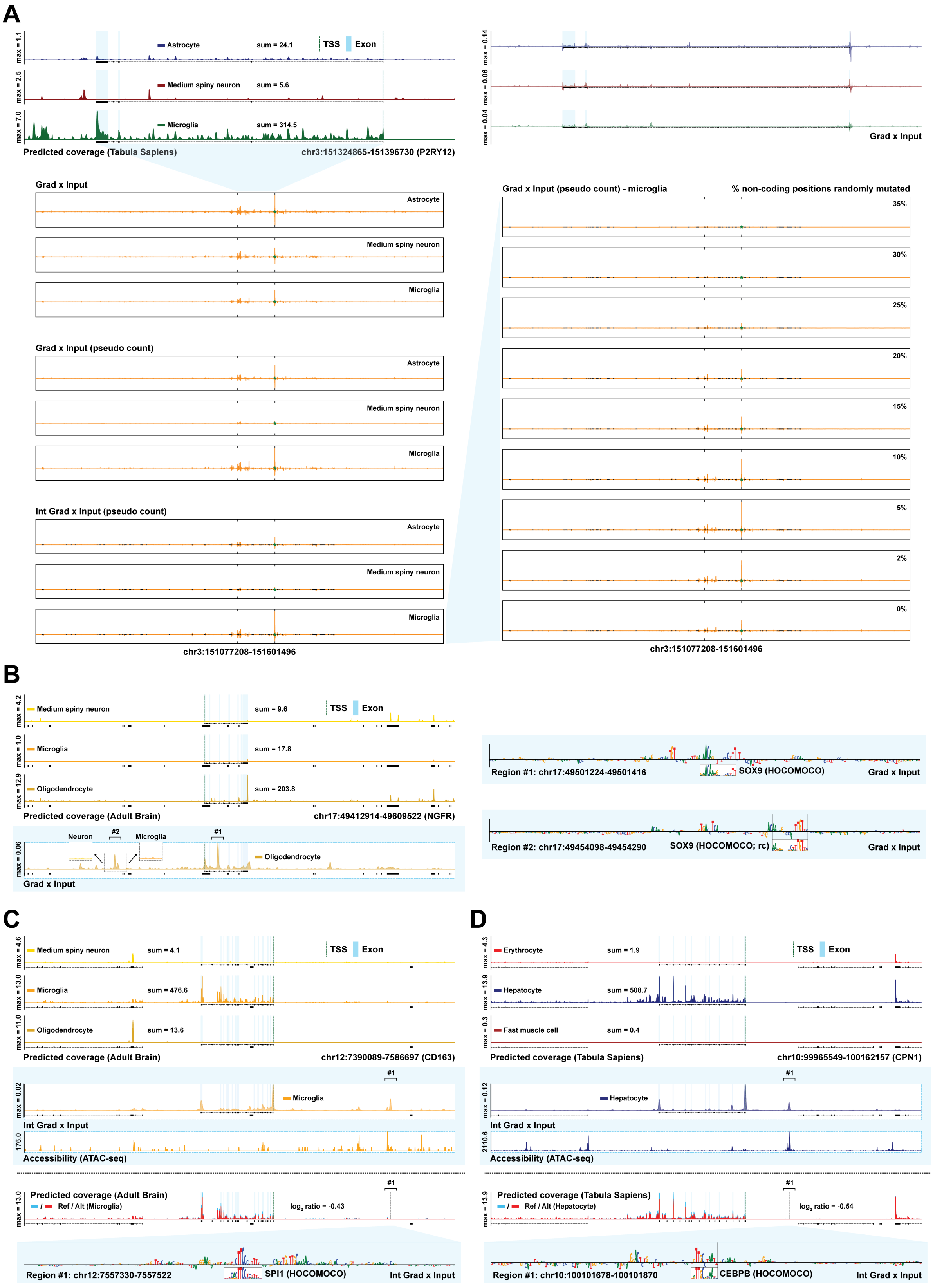
A
max = 1.1
Astrocyte
sum = 24.1
TSS
Exon
max = 2.5
Medium spiny neuron
sum = 5.6
max = 7.0
Microglia
sum = 314.5
Predicted coverage (Tabula Sapiens)
chr3:151324865-151396730 (P2RY12)
max = 0.14
max = 0.06
max = 0.04
Grad x Input
Grad x Input
Astrocyte
Medium spiny neuron
Microglia
Grad x Input (pseudo count)
Astrocyte
Medium spiny neuron
Microglia
Int Grad x Input (pseudo count)
Astrocyte
Medium spiny neuron
Microglia
chr3:151077208-151601496
Grad x Input (pseudo count) - microglia
% non-coding positions randomly mutated
35%
30%
25%
20%
15%
10%
5%
2%
0%
chr3:151077208-151601496
B
max = 4.2
Medium spiny neuron
sum = 9.6
TSS
Exon
max = 1.0
Microglia
sum = 17.8
max = 12.9
Oligodendrocyte
sum = 203.8
Predicted coverage (Adult Brain)
chr17:49412914-49609522 (NGFR)
max = 0.06
Neuron
#2
Microglia
#1
Oligodendrocyte
Grad x Input
Region #1: chr17:49501224-49501416
SOX9 (HOCOMOCO)
Grad x Input
Region #2: chr17:49454098-49454290
SOX9 (HOCOMOCO; rc)
Grad x Input
C
max = 4.6
Medium spiny neuron
sum = 4.1
TSS
Exon
max = 13.0
Microglia
sum = 476.6
max = 11.0
Oligodendrocyte
sum = 13.6
Predicted coverage (Adult Brain)
chr12:7390089-7586697 (CD163)
#1
max = 0.02
Microglia
Int Grad x Input
176.0
Accessibility (ATAC-seq)
#1
max = 13.0
Predicted coverage (Adult Brain)
Ref / Alt (Microglia)
log2 ratio = -0.43
Region #1: chr12:7557330-7557522
SPI1 (HOCOMOCO)
Int Grad x Input
D
max = 4.3
Erythrocyte
sum = 1.9
TSS
Exon
max = 13.9
Hepatocyte
sum = 508.7
max = 0.3
Fast muscle cell
sum = 0.4
Predicted coverage (Tabula Sapiens)
chr10:99965549-100162157 (CPN1)
#1
max = 0.12
Hepatocyte
Int Grad x Input
2110.6
Accessibility (ATAC-seq)
#1
max = 13.9
Predicted coverage (Tabula Sapiens)
Ref / Alt (Hepatocyte)
log2 ratio = -0.54
Region #1: chr10:100101678-100101870
CEBPB (HOCOMOCO)
Int Grad x Input

Supplementary Figure 2: **Related to Figure 2. (A)** Top: Visualization of predicted scRNA-seq coverage and input-gated gradient saliencies in three cell types from the Adult Brain atlas for gene *P2RY12*. The gradient saliencies are notably miscalibrated, with higher saliency at the promoter for Astrocyte than for Microglia. Bottom: Comparison of three different approaches for saliency estimation: (1) input-gated gradients, (2) input-gated gradients with a pseudo-count added to the predicted exon coverage before log-normalizing and computing the gradient, and (3) 'integrated' input-gated gradients, where the noncoding regions of the input sequence are randomly mutated according to a varying rate and gradients are re-computed and averaged for all mutant sequences. **(B)** Predicted scRNA-seq coverage for gene *NGFR*. Shown below are the standard gradient attributions of exon-aggregated coverage in Oligodendrocyte, with highlighted regions on the right (integrated gradients were displayed in Figure 2). **(C)** Predicted scRNA-seq coverage in three brain cell types for gene *CD163*, with (integrated) exon-aggregated coverage gradients shown for Microglia along with experimental ATAC-seq coverage. The bottom plot displays predicted coverage in Microglia after dinucleotide-shuffling the highlighted putative enhancer. **(D)** Predicted scRNA-seq coverage in three cell types from Tabula Sapiens for gene *CPN1*, with integrated gradient attributions and perturbation predictions shown below for Hepatocyte.

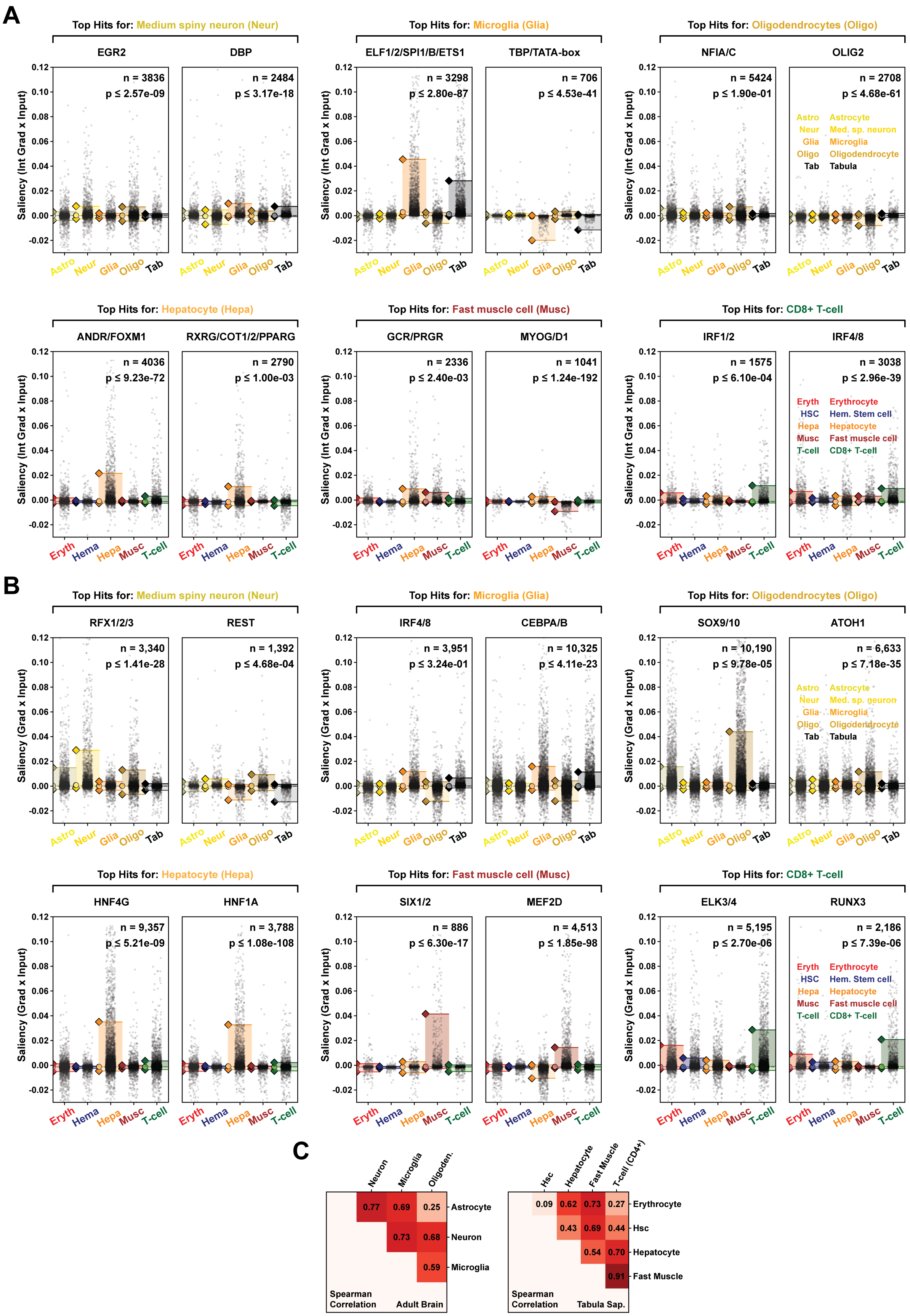
A
Top Hits for: Medium spiny neuron (Neur)
EGR2
n = 3836
p ≤ 2.57e-09
DBP
n = 2484
p ≤ 3.17e-18
Top Hits for: Microglia (Glia)
ELF1/2/SPI1/B/ETS1
n = 3298
p ≤ 2.80e-87
TBP/TATA-box
n = 706
p ≤ 4.53e-41
Top Hits for: Oligodendrocytes (Oligo)
NFIA/C
n = 5424
p ≤ 1.90e-01
OLIG2
n = 2708
p ≤ 4.68e-61
Astro Astrocyte
Neur Med. sp. neuron
Glia Microglia
Oligo Oligodendrocyte
Tab Tabula
Saliency (Int Grad x Input)
0.12
0.10
0.08
0.06
0.04
0.02
0.00
-0.02
Astro Neur Glia Oligo Tab
Top Hits for: Hepatocyte (Hepa)
ANDR/FOXM1
n = 4036
p ≤ 9.23e-72
RXRG/COT1/2/PPARG
n = 2790
p ≤ 1.00e-03
Top Hits for: Fast muscle cell (Musc)
GCR/PRGR
n = 2336
p ≤ 2.40e-03
MYOG/D1
n = 1041
p ≤ 1.24e-192
Top Hits for: CD8+ T-cell
IRF1/2
n = 1575
p ≤ 6.10e-04
IRF4/8
n = 3038
p ≤ 2.96e-39
Eryth Erythrocyte
HSC Hem. Stem cell
Hepa Hepatocyte
Musc Fast muscle cell
T-cell CD8+ T-cell
Eryth Hema Hepa Musc T-cell
B
Top Hits for: Medium spiny neuron (Neur)
RFX1/2/3
n = 3,340
p ≤ 1.41e-28
REST
n = 1,392
p ≤ 4.68e-04
Top Hits for: Microglia (Glia)
IRF4/8
n = 3,951
p ≤ 3.24e-01
CEBPA/B
n = 10,325
p ≤ 4.11e-23
Top Hits for: Oligodendrocytes (Oligo)
SOX9/10
n = 10,190
p ≤ 9.78e-05
ATOH1
n = 6,633
p ≤ 7.18e-35
Astro Astrocyte
Neur Med. sp. neuron
Glia Microglia
Oligo Oligodendrocyte
Tab Tabula
Saliency (Grad x Input)
Top Hits for: Hepatocyte (Hepa)
HNF4G
n = 9,357
p ≤ 5.21e-09
HNF1A
n = 3,788
p ≤ 1.08e-108
Top Hits for: Fast muscle cell (Musc)
SIX1/2
n = 886
p ≤ 6.30e-17
MEF2D
n = 4,513
p ≤ 1.85e-98
Top Hits for: CD8+ T-cell
ELK3/4
n = 5,195
p ≤ 2.70e-06
RUNX3
n = 2,186
p ≤ 7.39e-06
Eryth Erythrocyte
HSC Hem. Stem cell
Hepa Hepatocyte
Musc Fast muscle cell
T-cell CD8+ T-cell
C
Neuron Microglia Oligoden.
0.77 0.69 0.25 Astrocyte
0.73 0.68 Neuron
0.59 Microglia
Spearman Correlation
Adult Brain
Hsc Hepatocyte Fast Muscle T-cell (CD4+)
0.09 0.62 0.73 0.27 Erythrocyte
0.43 0.69 0.44 Hsc
0.54 0.70 Hepatocyte
0.91 Fast Muscle
Spearman Correlation
Tabula Sap.

Supplementary Figure 3: **Related to Figure 2. (A)** Integrated gradient saliency distributions for an additional set of putative TF motifs. The top row displays putative motifs identified for cell types from the Adult Brain atlas, and the bottom row displays motifs for Tabula Sapiens cell types. Saliencies were calculated with respect to the predicted exon-aggregated log-coverage of $1,000$ differentially expressed genes for the Adult Brain and Tabula Sapiens. Each dot in the distribution plots represent a unique genomic instance of a given motif. P-values were computed as two-sided Wilcoxon tests between the cell types with largest and second largest 95th percentile of absolute-valued saliencies. **(B)** Re-visualizing saliency distributions for the same selection of putative motifs as shown in Figure 2, but where the saliencies are derived from standard input gradients rather than integrated gradients. **(C)** Spearman correlation comparing the average difference in motif saliency for pairs of cell types to the average difference in log-TPM for the corresponding TF gene. The TPM values of TF genes belonging to the same HOCOMOCO subfamily were averaged. The sign of the saliency scores for REST, OLIG2, and MyoD was inverted to reflect their well-established repressive roles.

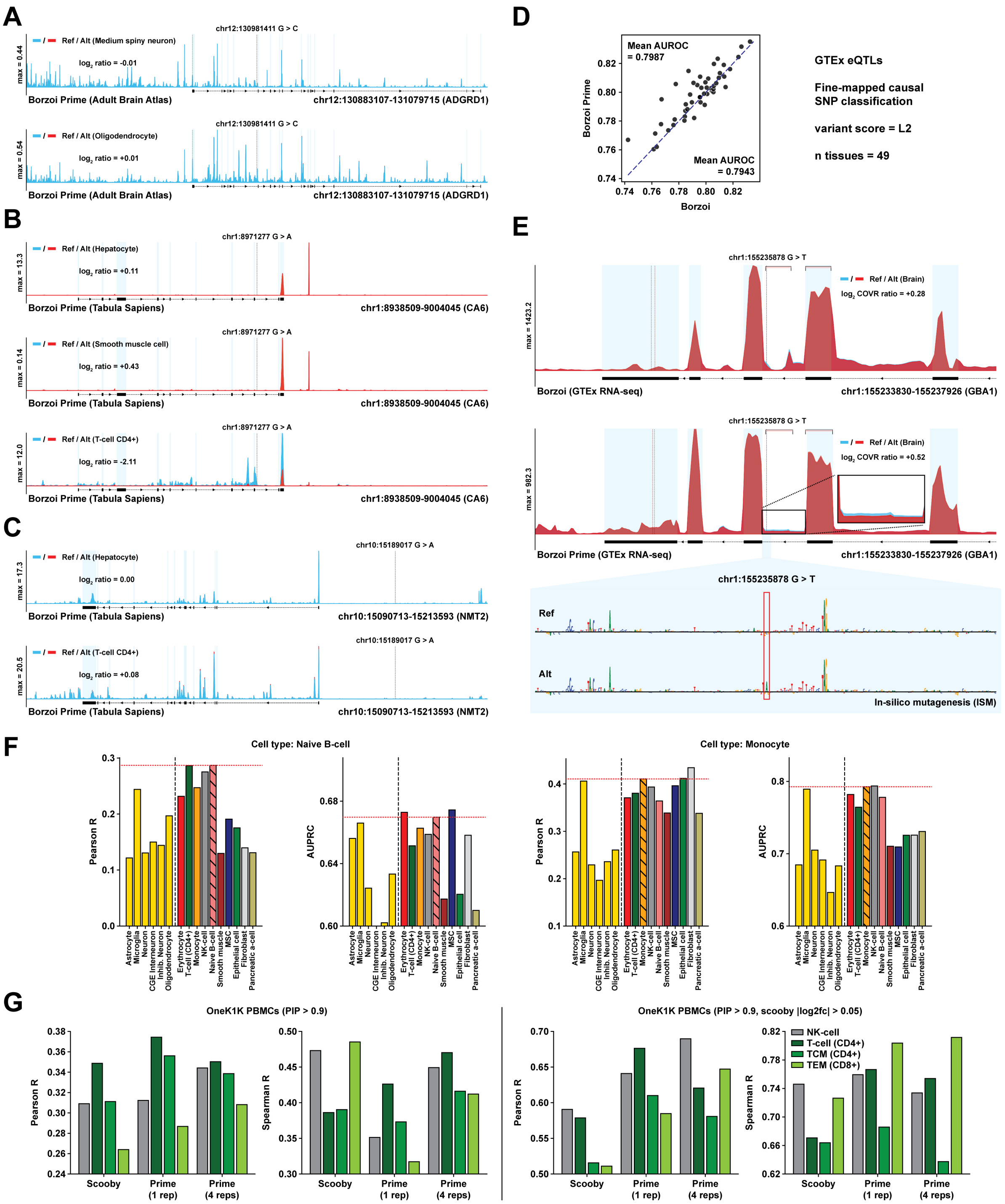
A
chr12:130981411 G > C
Ref / Alt (Medium spiny neuron)
log2 ratio = -0.01
max = 0.44
Borzoi Prime (Adult Brain Atlas)
chr12:130883107-131079715 (ADGRD1)
Ref / Alt (Oligodendrocyte)
log2 ratio = +0.01
max = 0.54
B
chr1:8971277 G > A
Ref / Alt (Hepatocyte)
log2 ratio = +0.11
max = 13.3
Borzoi Prime (Tabula Sapiens)
chr1:8938509-9004045 (CA6)
Ref / Alt (Smooth muscle cell)
log2 ratio = +0.43
max = 0.14
Ref / Alt (T-cell CD4+)
log2 ratio = -2.11
max = 12.0
C
chr10:15189017 G > A
Ref / Alt (Hepatocyte)
log2 ratio = 0.00
max = 17.3
chr10:15090713-15213593 (NMT2)
Ref / Alt (T-cell CD4+)
log2 ratio = +0.08
max = 20.5
D
Mean AUROC = 0.7987
Mean AUROC = 0.7943
Borzoi Prime
Borzoi
GTEx eQTLs
Fine-mapped causal SNP classification
variant score = L2
n tissues = 49
E
chr1:155235878 G > T
Ref / Alt (Brain)
log2 COVR ratio = +0.28
max = 1423.2
Borzoi (GTEx RNA-seq)
chr1:155233830-155237926 (GBA1)
log2 COVR ratio = +0.52
max = 982.3
Borzoi Prime (GTEx RNA-seq)
Ref
Alt
In-silico mutagenesis (ISM)
F
Cell type: Naive B-cell
Cell type: Monocyte
Pearson R
AUPRC
Astrocyte
Microglia
Neuron
CGE Interneuron
Inhib. Neuron
Oligodendrocyte
Erythrocyte
T-cell (CD4+)
Monocyte
NK-cell
Naive B-cell
Smooth muscle
MSC
Epithelial cell
Fibroblast
Pancreatic a-cell
G
OneK1K PBMCs (PIP > 0.9)
OneK1K PBMCs (PIP > 0.9, scooby |log2fc| > 0.05)
Pearson R
Spearman R
Scooby
Prime (1 rep)
Prime (4 reps)
NK-cell
T-cell (CD4+)
TCM (CD4+)
TEM (CD8+)

Supplementary Figure 4: **Related to Figure 3.** **(A)** Predicted scRNA-seq coverage for the wildtype sequence (blue) and mutated sequence (red) when substituting the alternate allele of fine-mapped eQTL rs12821379, shown in neuronal- and oligodendrocyte cell types. **(B)** Predicted scRNA-seq coverage in three Tabula Sapiens cell types for fine-mapped eQTL SNP rs116366173, showing reference (blue) and alternate (red) predictions. The SNP has a significant measured association in CD4+ T-cells. **(C)** Predicted scRNA-seq coverage in two Tabula Sapiens cell types for fine-mapped eQTL SNP rs76459781 (significant association in CD4+ T-cells). **(D)** Comparison between Borzoi Prime and Borzoi at classifying fine-mapped GTEx eQTL SNPs from matched negatives based on gene-agnostic variant scores derived from the L2-norm between reference- and alternate coverage predictions. An auxiliary random forest classifier is trained on the full set of tissue- and cell type-specific L2-scores predicted by each model and used to classify negative from positive SNPs in a cross-validation setting. **(E)** Predicted GTEx bulk RNA-seq coverage in brain tissue for SNP rs3115534, showing reference predictions in blue and predictions of the mutated sequence in red for both Borzoi and Borzoi Prime. **(F)** Performance metrics when using Borzoi Prime to score sc-eQTLs for two additional cell types from the OneK1K study—Naive B-cells and Monocytes. Variant predictions were derived from cell type-matched scRNA-seq tracks from Tabula Sapiens, using track-specific pseudo-counts added to the reference- and alternate predicted coverage values before computing their log-ratio. The pseudo-count was calculated as the 20th percentile of gene expression values for each track. **(G)** Comparison between Borzoi Prime and scooby (fine-tuned on OneK1K multiome data) at predicting fine-mapped PBMC sc-eQTL effect sizes. Comparisons are shown for $4$ cell types with $> 100$ fine-mapped SNPs per cell type. Both Spearman and Pearson correlation coefficients are displayed. Left: All SNPs with PIP $> 0.90$. Right: Subset of high-scoring SNPs. For scooby, only SNPs with a predicted absolute-valued fold change $> 3.5\%$ are shown (threshold obtained from scooby paper). For Borzoi Prime, we kept a matching number of SNPs, sorted in descending order of absolute-valued predicted effect.

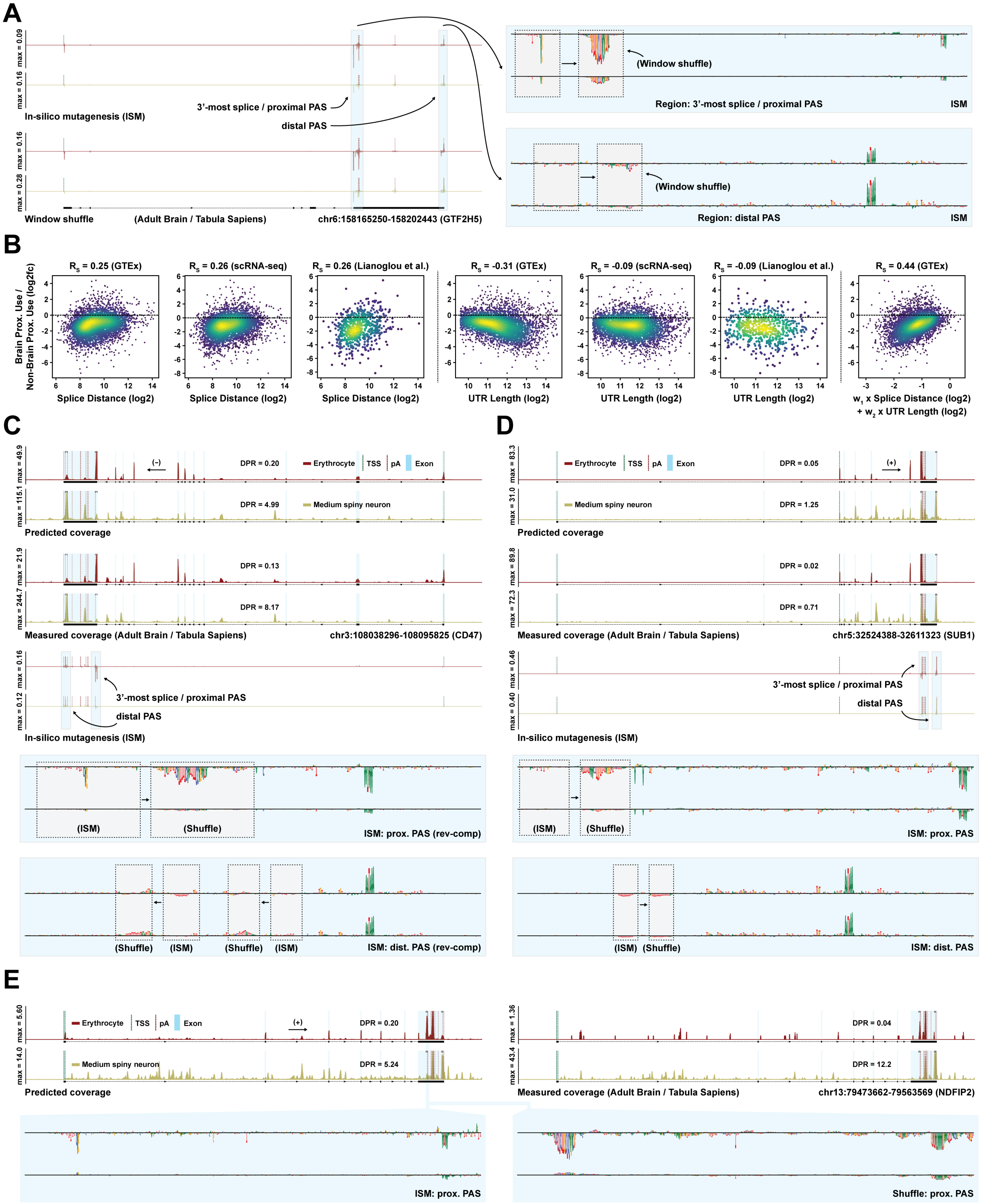
A
In-silico mutagenesis (ISM)
3'-most splice / proximal PAS
distal PAS
Window shuffle
(Adult Brain / Tabula Sapiens)
chr6:158165250-158202443 (GTF2H5)
(Window shuffle)
Region: 3'-most splice / proximal PAS
ISM
Region: distal PAS
B
Rs = 0.25 (GTEx)
Rs = 0.26 (scRNA-seq)
Rs = 0.26 (Lianoglou et al.)
Rs = -0.31 (GTEx)
Rs = -0.09 (scRNA-seq)
Rs = -0.09 (Lianoglou et al.)
Rs = 0.44 (GTEx)
Brain Prox. Use / Non-Brain Prox. Use (log2fc)
Splice Distance (log2)
UTR Length (log2)
w1 x Splice Distance (log2) + w2 x UTR Length (log2)
C
Erythrocyte
TSS
pA
Exon
Medium spiny neuron
DPR = 0.20
DPR = 4.99
DPR = 0.13
DPR = 8.17
Predicted coverage
Measured coverage (Adult Brain / Tabula Sapiens)
chr3:108038296-108095825 (CD47)
(ISM)
(Shuffle)
ISM: prox. PAS (rev-comp)
ISM: dist. PAS (rev-comp)
D
DPR = 0.05
DPR = 1.25
DPR = 0.02
DPR = 0.71
chr5:32524388-32611323 (SUB1)
ISM: prox. PAS
ISM: dist. PAS
E
DPR = 0.20
DPR = 5.24
DPR = 0.04
DPR = 12.2
chr13:79473662-79563569 (NDFIP2)
Shuffle: prox. PAS

Supplementary Figure 5: **Related to Figure 4.** **(A)** In-silico saturation mutagenesis (ISM) and window-shuffled sequence attributions in 512bp windows around each annotated 3' UTR polyadenylation site or splice junction of the *GTF2H5* gene. The attributions are calculated with respect to predicted distal polyadenylation site usage and are shown for Erythrocytes and Medium spiny neurons. Right: Saliency scores in regions overlapping the proximal-most and distal-most polyadenylation site, displayed as sequence logos. All attribution tracks and sequence logos are shown with equally scaled y-axis. **(B)** Comparison between various features extracted from GENCODE v41 annotations (x-axis) and the gene-specific proximal polyadenylation bias in brain relative to non-brain measurements (y-axis). The annotated features include (1) $\log_2$-distance between the 3'-most splice acceptor and proximal-most polyadenylation site, (2) $\log_2$-length of the 3' UTR, or (3) linear combination of these two features (learned from linear regression with 10-fold cross-validation). The measurements displayed along the y-axis vary depending on source; for GTEx, we calculate the $\log_2$-fold change between the proximal coverage ratio in brain and the average proximal coverage ratio in blood or adipose tissue. For scRNA-seq, we calculate this fold change between pseudo-bulked coverage ratios from the Adult Brain atlas and blood- or adipose tissue from Tabula Sapiens. For bulk 3'-seq data curated from PolyASite 2.0, we calculated fold changes in proximal usage between brain- and B-LCL samples from Lianoglou et al. (2013) (SRA identifiers: SRX359339, SRX351952). **(C)** Predicted and measured scRNA-seq coverage in Erythrocytes and Medium spiny neurons for gene *CD47*. ISM sequence attributions for annotated splice junctions and polyadenylation signals are shown below, with sequence logos displayed for the proximal-most and distal-most polyadenylation signals. Window-shuffled attributions for putative cell type-specific motif regions are displayed as insets in the sequence logos. DPR = ratio of distal-to-proximal coverage. **(D)** Coverage predictions, measurements, and ISM attributions for gene *SUB1*. **(E)** Coverage predictions, measurements, and ISM attributions for gene *NDFIP2*.

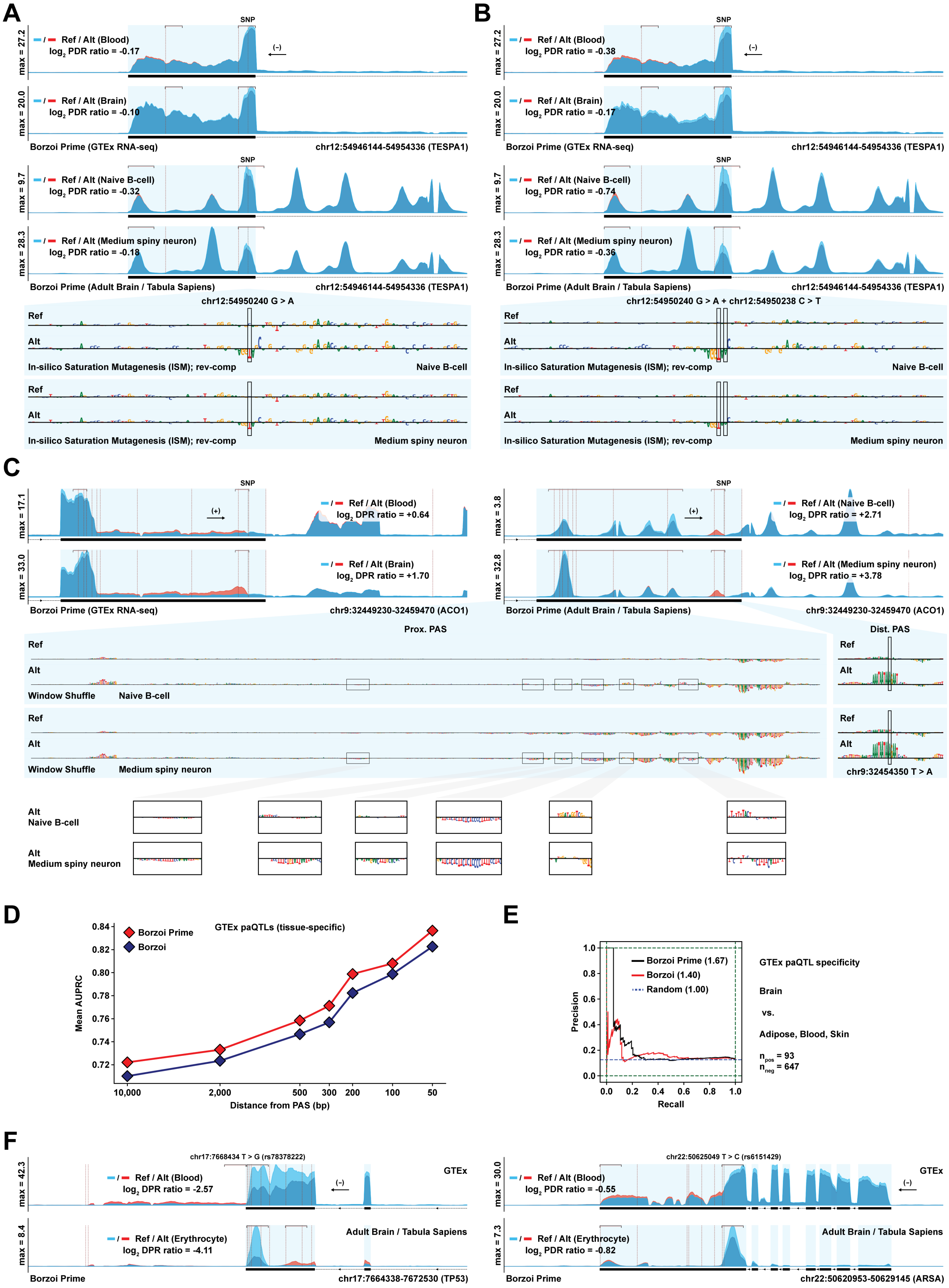
A
SNP
(−)
Ref / Alt (Blood)
log2 PDR ratio = -0.17
Ref / Alt (Brain)
log2 PDR ratio = -0.10
max = 27.2
max = 20.0
Borzoi Prime (GTEx RNA-seq)
chr12:54946144-54954336 (TESPA1)
Ref / Alt (Naive B-cell)
log2 PDR ratio = -0.32
Ref / Alt (Medium spiny neuron)
log2 PDR ratio = -0.18
max = 9.7
max = 28.3
Borzoi Prime (Adult Brain / Tabula Sapiens)
chr12:54946144-54954336 (TESPA1)
chr12:54950240 G > A
Ref
Alt
In-silico Saturation Mutagenesis (ISM); rev-comp
Naive B-cell
Medium spiny neuron
B
Ref / Alt (Blood)
log2 PDR ratio = -0.38
Ref / Alt (Brain)
log2 PDR ratio = -0.17
Ref / Alt (Naive B-cell)
log2 PDR ratio = -0.74
Ref / Alt (Medium spiny neuron)
log2 PDR ratio = -0.36
chr12:54950240 G > A + chr12:54950238 C > T
C
(+)
Ref / Alt (Blood)
log2 DPR ratio = +0.64
Ref / Alt (Brain)
log2 DPR ratio = +1.70
max = 17.1
max = 33.0
Borzoi Prime (GTEx RNA-seq)
chr9:32449230-32459470 (ACO1)
Ref / Alt (Naive B-cell)
log2 DPR ratio = +2.71
Ref / Alt (Medium spiny neuron)
log2 DPR ratio = +3.78
max = 3.8
max = 32.8
Borzoi Prime (Adult Brain / Tabula Sapiens)
chr9:32449230-32459470 (ACO1)
Prox. PAS
Dist. PAS
Window Shuffle
Naive B-cell
Medium spiny neuron
chr9:32454350 T > A
Alt
Naive B-cell
Alt
Medium spiny neuron
D
GTEx paQTLs (tissue-specific)
Borzoi Prime
Borzoi
Mean AUPRC
0.84
0.82
0.80
0.78
0.76
0.74
0.72
10,000
2,000
500
300
200
100
50
Distance from PAS (bp)
E
Borzoi Prime (1.67)
Borzoi (1.40)
Random (1.00)
Precision
Recall
1.0
0.8
0.6
0.4
0.2
0.0
GTEx paQTL specificity
Brain
vs.
Adipose, Blood, Skin
npos = 93
nneg = 647
F
chr17:7668434 T > G (rs78378222)
GTEx
Ref / Alt (Blood)
log2 DPR ratio = -2.57
Ref / Alt (Erythrocyte)
log2 DPR ratio = -4.11
max = 42.3
max = 8.4
Adult Brain / Tabula Sapiens
Borzoi Prime
chr17:7664338-7672530 (TP53)
chr22:50625049 T > C (rs6151429)
Ref / Alt (Blood)
log2 DPR ratio = -0.55
Ref / Alt (Erythrocyte)
log2 DPR ratio = -0.82
max = 30.0
max = 7.3
chr22:50620953-50629145 (ARSA)

Supplementary Figure 6: **Related to Figure 4.** **(A)** Predicted bulk RNA-seq coverage in GTEx tissues blood and brain and scRNA-seq coverage in naive B-cells and medium spiny neurons for the fine-mapped eQTL SNP rs1801876 in the gene *TESPA1*. Shown are the predictions for the reference (blue) and alternative (red) sequences. ISM attributions are displayed for B-cells and neurons at the bottom, indicating a gain of a non-canonical U1 snRNA motif due to the variant. PDR = ratio of proximal-to-distal coverage. **(B)** Predicted coverage for a hypothetical double mutation (not found in the population) in the *TESPA1* gene, creating a canonical U1 snRNA motif and resulting in a more pronounced cell type-specific functional impact. **(C)** Reference- and variant predictions for fine-mapped eQTL SNP rs1360171. The tracks display predicted GTEx RNA-seq coverage in blood and brain and scRNA-seq coverage in naive B-cells and neurons. Sequence attributions based on window-shuffled ISM in B-cells and neurons are shown at the bottom. While no cell type-specific features are found in the distal site near the mutation, there are a number of putative cell type-specific motifs identified in the saliency scores around the proximal site (U- and UC-rich motifs, putative NOVA1 binding sites, etc.). DPR = ratio of distal-to-proximal coverage. **(D)** Comparison between Borzoi Prime and Borzoi at distinguishing fine-mapped GTEx paQTL SNPs and distance-matched negatives. A tissue-pooled version of this analysis is presented in Figure 4, where the classification AUPRC is calculated for the union of positive and negative SNPs across all tissues. Here, we instead report the average tissue-specific AUPRC metric, plotted as a function of the maximum distance to the nearest annotated polyadenylation site. **(E)** Comparison between Borzoi Prime and Borzoi when predicting whether a fine-mapped causal paQTL has a cell type-specific effect in brain or whether it has a specific or shared effect in adipose, skin, or blood. Shown are the precision-recall curves for both models, as well as the theoretical precision of a random predictor. AUPRC metrics are annotated in the legend. **(F)** Variant effect predictions based on GTEx blood RNA-seq tracks for two known pathogenic mutations: rs78378222 (TP53) and rs6151429 (ARSA). Shown are the predicted coverage for the reference (blue) and alternative (red) sequences.
